# Filopodome mapping identifies p130Cas as a mechanosensitive regulator of filopodia stability

**DOI:** 10.1101/377879

**Authors:** Guillaume Jacquemet, Rafael Saup, Hellyeh Hamidi, Mitro Miihkinen, Johanna Ivaska

## Abstract

Filopodia are adhesive cellular protrusions specialised in the detection of extracellular matrix (ECM)-derived cues. While ECM engagement at focal adhesions is known to trigger the recruitment of hundreds of proteins (“adhesome”) to fine-tune cellular behaviour, the components of the filopodia adhesions remain undefined. Here, we performed a structured illumination microscopy-based screen to map the localisation of 80 target proteins, linked to cell adhesion and migration, within filopodia. We demonstrate preferential enrichment of several adhesion proteins to either filopodia tips, filopodia shafts, or shaft subdomains suggesting divergent, spatially restricted functions for these proteins. Moreover, proteins with phosphoinositide (PI) binding sites are particularly enriched in filopodia. This, together with the strong localisation of PI(3,4)P_2_ in filopodia tips, predicts critical roles for PIs in regulating filopodia ultra-structure and function. Our mapping further reveals that filopodia adhesions consist of a unique set of proteins, the filopodome, that are distinct from classical nascent adhesions, focal adhesions and fibrillar adhesions. Using live imaging, we observe that filopodia adhesions can give rise to nascent adhesions, which, in turn, form focal adhesions. Finally, we demonstrate that p130Cas (BCAR1) is recruited to filopodia tips via its CCHD domain and acts as a mechanosensitive regulator of filopodia stability.

## Introduction

The ability of cells to migrate in vivo is necessary for many physiological processes including embryonic development, tissue homoeostasis and wound healing. Cell migration is also implicated in distinct pathological conditions such as inflammation and cancer metastasis. To migrate, cells interact with their environment, the extracellular matrix (ECM), via adhesion receptors such as integrins which provide a physical link between the ECM and the actin cytoskeleton (Legate et al., 2009). Integrin function is controlled by a conformational switch between active and inactive states that determine ECM ligand interaction and subsequent receptor signalling (Askari et al., 2009). Integrin activation can be triggered from within the cell by several mechanisms including the Rap1-RIAM-talin pathway. In 2D, integrin-ligand engagement leads to the assembly of large signalling platforms, termed focal adhesions (FA), which are composed of hundreds of proteins collectively termed the adhesome (Horton et al., 2016; Zaidel-Bar and Geiger, 2010; Zaidel-Bar et al., 2007). FA are highly dynamic and complex structures that develop from force-dependent maturation of nascent adhesions at the leading edge, and which undergo integrin-specific centripetal translocation to form fibrillar adhesions. Importantly FA not only provide anchorage but also represent integrin heterodimer-ligand-specific (Morgan et al., 2009) and/or ECM-ligand-specific signalling nodes with mechanosensing functions (Jansen et al., 2017), and, therefore constitute ideal signalling platforms for ECM recognition.

Cell motility through complex 3D microenvironments also requires efficient probing of the cell surroundings via specialized sensory protrusions such as filopodia (Jacquemet et al., 2015). Filopodia are finger-like actin-rich protrusions widely used by cells in vivo during normal processes, such as development (Sato et al., 2017; Haas and Gilmour, 2006; Zhang et al., 2018), angiogenesis (Gerhardt et al., 2003), immune surveillance (Liu et al., 2018), and wound healing (Wood et al., 2002; Pickering et al., 2013), and also during tumourigenesis (Shibue et al., 2012, 2013) and cancer cell dissemination (Paul et al., 2015; Jacquemet et al., 2017; Follain et al., 2018; Liu et al., 2018). Filopodia have integrin-positive adhesive tips for ECM sensing and cells can employ filopodia to recognize ECM gradients (Johnson et al., 2015), ECM topography (Albuschies and Vogel, 2013) or ECM stiffness (Wong et al., 2014; Chan and Odde, 2008). Thus, filopodia contribute to haptotaxis (Johnson et al., 2015) and possibly to durotaxis (Kim et al., 2018). Extension of a filopodium is driven by linear polymerization and bundling of actin filaments mediated by proteins such as formins and fascin, respectively (Heckman and Plummer, 2013). Filopodia are very dynamic structures that stabilise upon ECM tethering and formation of an adhesive structure, at their tips, termed filopodia adhesion. ECM attachment at filopodia is mediated by integrins, which are actively transported to filopodia tips by the motor protein myosin-X (MYO10) (Berg and Cheney, 2002; Zhang et al., 2004). In addition, filopodia stabilisation requires both talin-mediated integrin activation and integrin downstream signalling at the filopodium tip (Jacquemet et al., 2016). However, the composition of filopodia adhesions remains poorly defined.

The composition and architecture of FA have been extensively studied using both microscopy and mass spectrometric-based strategies (Kanchanawong et al., 2010; Horton et al., 2015; Zaidel-Bar and Geiger, 2010). Importantly the compilation of FA components, either by literature curation (Geiger Adhesome, (Zaidel-Bar and Geiger, 2010; Zaidel-Bar et al., 2007)) or by proteomic approaches (Consensus Adhesome, (Horton et al., 2015, 2016)), have led to significant advances in our understanding of adhesion-mediated processes. Considering that filopodia are significant structures in vivo that are also implicated in various biological processes and pathologies such as angiogenesis and cancer progression (Jacquemet et al., 2017), a more detailed analysis of proteins recruited to filopodia adhesions is likely to be fundamental to better understand filopodia functions. Filopodia are relatively small and labile structures (1-5 μm length and 50-200 nm width) and are therefore difficult to purify in a scale sufficient to perform mass-spectrometry analyses. Here, to characterize the composition of filopodia tip adhesions, we used a targeted approach and mapped the localisation of 80 proteins, implicated in cellular adhesion or protein-membrane lipid interactions, using structured illumination microscopy (SIM).

## Results

### Mapping protein localisation in filopodia using structured illumination microscopy

Filopodia are relatively small structures that are difficult to image using diffraction limited approaches. We found that SIM imaging overcomes this technical challenge and is an ideal strategy to simultaneously visualise the localisation of up to three proteins within filopodia at high resolution (Figure 1A and 1B). To identify proteins that localise to filopodia, we performed a SIM-based screen, spanning 80 putative regulators of cellular adhesion or protein-membrane lipid association. Proteins of interests (POI, Figure 1C and Table S1) included known filopodia components, actin regulators, established FA components (Geiger Adhesome, (Zaidel-Bar and Geiger, 2010; Zaidel-Bar et al., 2007)) and adhesion proteins consistently identified in multiple mass spectrometry studies (Consensus Adhesome, (Horton et al., 2015)). We chose to visualize all the POI as GFP-fusion proteins, as it was not feasible to generate, validate and optimize antibodies against all endogenous POI. Cells adhering to fibronectin and transiently co-expressing a GFP-tagged POI and RFP-MYO10 (to visualise filopodia tips) were stained for F-actin and imaged using SIM (Figure 1A). SIM images of each POI are provided as supplementary information (Table S1, Figure S1-S11). We first observed the FA accumulation of each POI (Figure 1C and Figure S1-S11). Strikingly, 15 of the 38 Consensus Adhesome proteins (Horton et al., 2015) imaged did not display clear accumulation in FA (Figure 1C and Figure S1-S11), suggesting that they may be involved in other types of cell-ECM interface structures. To study the localisation of each POI (Table S1) to filopodia and their distribution along filopodia, line intensity profiles, manually drawn from filopodium tip to base (Figure 1A and 1B), were obtained for more than two hundred filopodia per POI. Importantly, to compare the distribution of POI in multiple cells, the brightness/contrast of each image was automatically adjusted using, as an upper maximum, the brightest cellular structure labelled. Based on line intensity profiles, the percentage of filopodia positive for each POI was quantified (Figure 1C) and a map highlighting the distribution of the POI within filopodia was created (Figure 2, see methods for details). The ImageJ macro and R scripts used to perform these quantifications are available as supplementary files (Scripts 1-3).

**Fig. 1.**
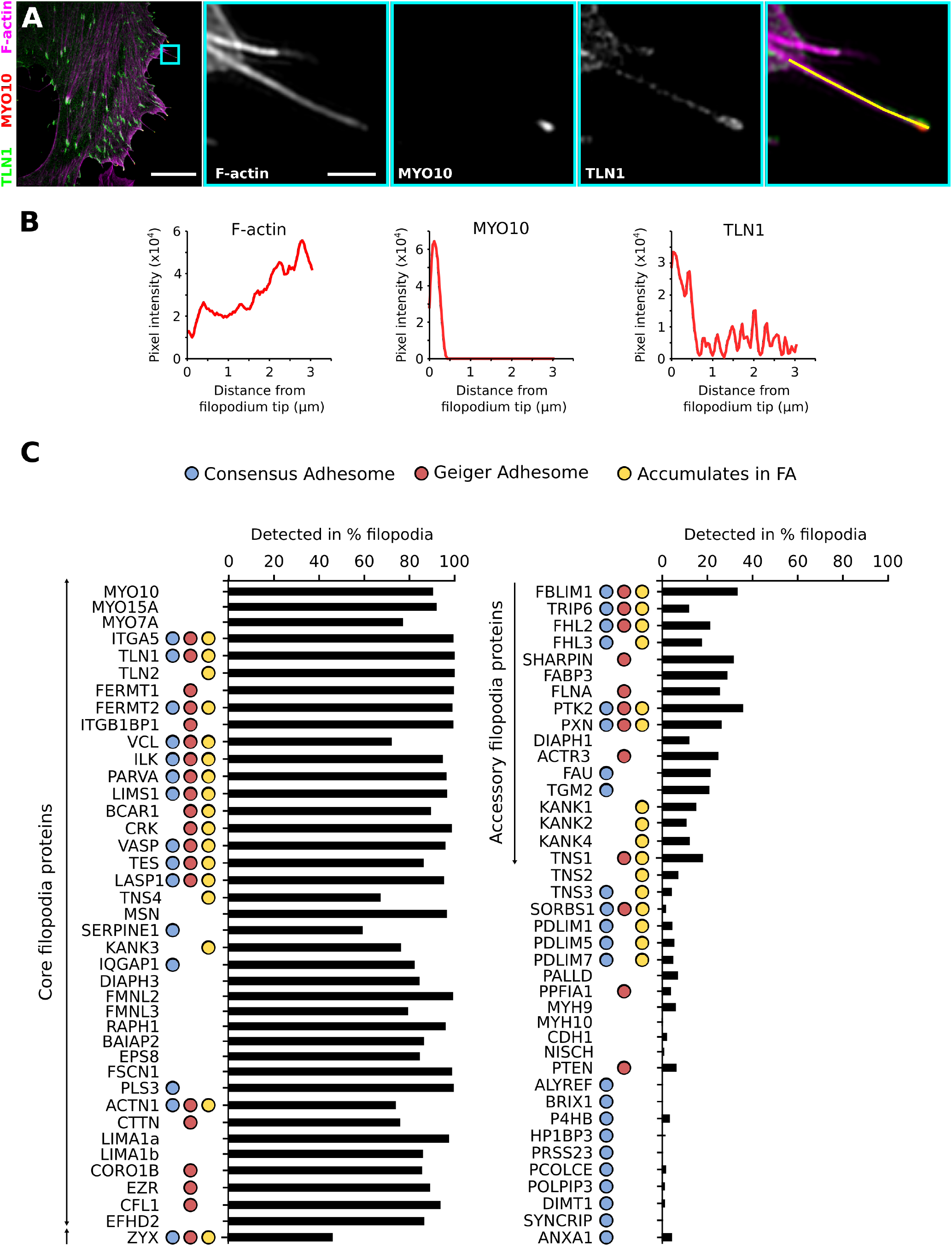
Mapping adhesion proteins to filopodia using structured illumination microscopy (SIM). A-C: To map the localisation of 80 key adhesion proteins within filopodia, U2OS cells expressing a GFP/RFP-tagged protein of interest (POI) and GFP/RFP-MYO10 were plated on fibronectin for 2 h, fixed and stained for F-actin before being imaged using SIM. To assess the distribution of each POI within filopodia, line intensity profiles were obtained, from filopodium tip to base, across hundreds of filopodia. A: An example illustrating the distribution of GFP-TLN1 (talin-1), MYO10-mScarlet (myosin-X) and F-actin within filopodia is shown. The blue square highlights the region of interest (ROI) that is magnified. Scale bars: (main) 10 μm; (inset) 1 μm. The yellow line represents the line drawn to measure the TLN1, MYO10 and F-actin intensity profiles. B: Results of the line intensity profiles from A. C: Results from line intensity profiles were used to quantify the percentage of filopodia positive for each POI (see methods for details). Each POI is designated by its official human gene name. Next to each POI, coloured circles indicate if this POI belongs to the Geiger Adhesome ((Zaidel-Bar and Geiger, 2010; Zaidel-Bar et al., 2007), red circle), to the Consensus Adhesome ((Horton et al., 2015), blue circle) or if this POI was found to accumulate in focal adhesions (FA) (this study, yellow circle). “Core filopodia proteins” and “accessory filopodia proteins” are labelled.

**Fig. 2.**
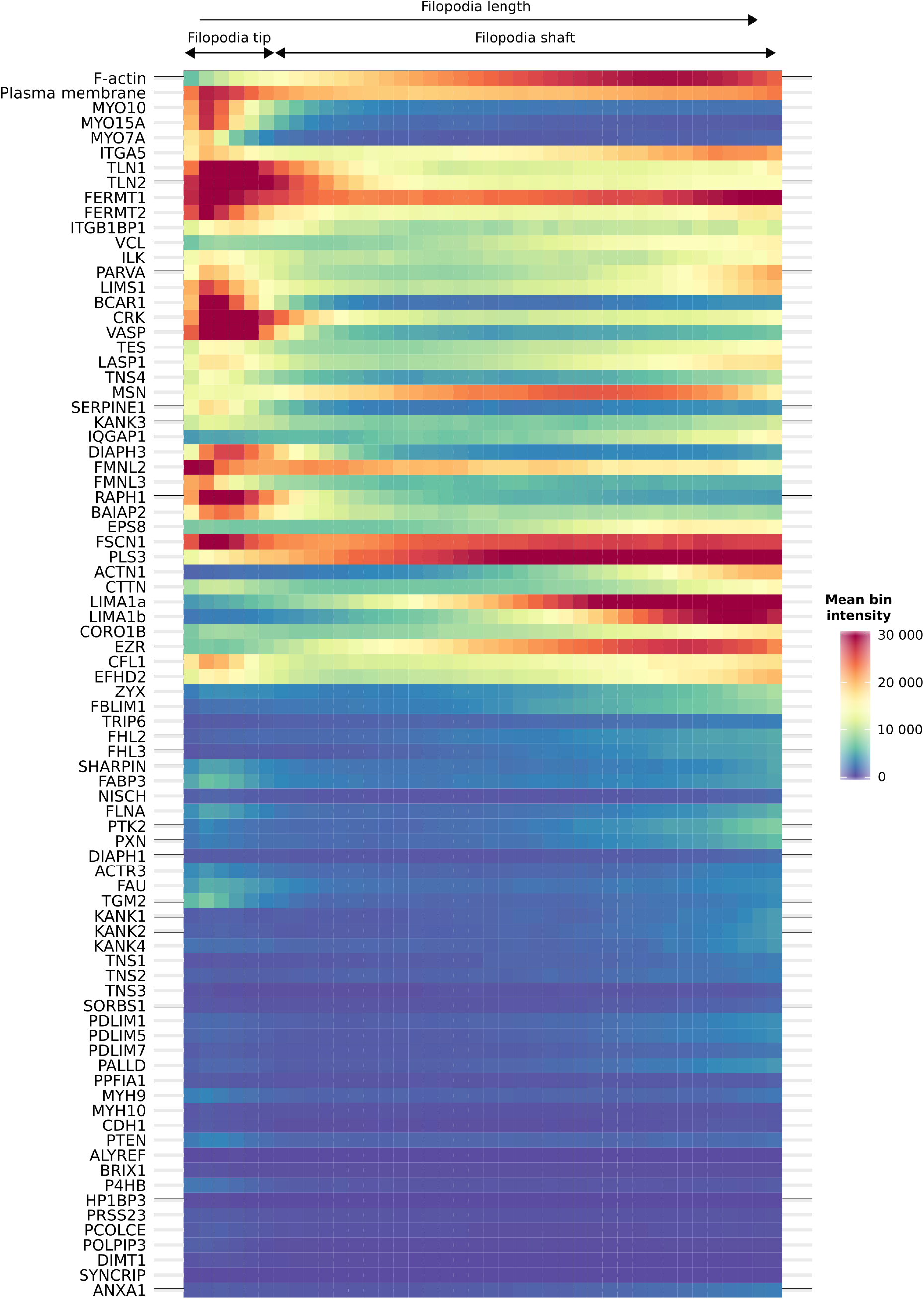
Generation of a filopodia map using correlative imaging and intensity profile averaging. Mapping of the subcellular localisation of 80 adhesion proteins (POI) within filopodia using at least 250 intensity profiles per POI (measured as in Figure 1; n numbers can be found in Table S1). POI are labelled using their official human gene name. To create this map, intensity profiles of each POI were binned (40 bins per filopodium / intensity profile) and each bin was then averaged (see methods for more details). The averaged intensity of each bin is colour-coded and the filopodia distribution of the proteins displayed as a heatmap. Location of the filopodium tip (defined by MYO10) and of the filopodium shaft are highlighted.

### Protein mapping reveals classes of core and accessory filopodia proteins

Multiple established filopodia-localising proteins including talin-1 (TLN1, (Lagarrigue et al., 2015)), formin-like protein 3 (FMNL3 (Harris et al., 2010)), lamellipodin (RAPH1, (Krause et al., 2004)), vasodilator-stimulated phosphoprotein (VASP; (Rottner et al., 1999)), mDia2 (DIAPH3, (Yang et al., 2007)) and fascin (FSCN1, (Vignjevic et al., 2006)) were clearly detected with high resolution in filopodia, validating our approach. Our comprehensive mapping revealed that the proteins imaged here can be organised into three categories in function of the frequency of their detection in filopodia. The first category is composed of the POI that are primarily absent from filopodia (Table S1, Figure 1C) and due to the low detection rate we consider that these proteins are not filopodia proteins (or are present in filopodia at very low density). The second category of POI are detected in a high proportion of filopodia (60% - 100%) indicating that these proteins are core filopodia proteins (Table S1, Figure 1C). The third category of POI are those reliably detected but present in only a small fraction of filopodia (10% - 40%) suggesting that these proteins may be accessory filopodia proteins contributing to filopodia-specific functions or defining subsets of biologically distinct filopodia (Table S1, Figure 1C).

### Phosphoinositide PI(3,4)P_2_ is enriched at filopodia tips

To identify common elements among the core filopodia proteins identified here, a protein domain enrichment analysis was performed (Figure 3A). This analysis revealed a strong enrichment of proteins containing FERM (24%) and/or pleckstrin homology (PH)-like (34%) and/or SRC homology 3 (SH3) domains (21%) among the proteins localising to filopodia (Figure 3A). A strong enrichment of proteins containing PH-like domains led us to speculate that the phosphoinositide (PI) composition of filopodia could be a key contributor to filopodia function. PI are major membrane-bound signalling molecules that, in function of their phosphorylation status, trigger the recruitment of adaptors to the plasma membrane to trigger signalling cascades (Bunney and Katan, 2010). To map the distribution of the various PI in filopodia, GFP-tagged probes with high affinity to a single PI species (Maekawa and Fairn, 2014; Idevall-Hagren and Camilli, 2015) were imaged using SIM and analysed as described above (Figure 3B and 3C). PI(3)P (imaged using GFP-FYVE-PH) was mostly detected on vesicular structures within the cell body, as previously reported (Idevall-Hagren and Camilli, 2015), but not in filopodia (Figure 3B and 3C). PI(4)P (imaged using GFP-P4M, (Hammond et al., 2014)) localised at the plasma membrane and on intracellular vesicles but was only weakly detected within filopodia (Figure 3B and 3C). In contrast, PI(4,5)P_2_ (labelled with GFP-PLC_γ_PH) and PI(3,4,5)P_3_ (labelled with GFP-BTK-PH) were both strongly detected within filopodia where their distribution was relatively homogeneous (similar to the plasma membrane probe distribution). Strikingly PI(3,4)P_2_(labelled using GFP-TAAP-PH) was also detected in filopodia, but in contrast to the other PI, PI(3,4)P_2_ was strongly enriched to filopodia tips. This strong accumulation of PI(3,4)P_2_ to filopodia tips is surprising and suggests that PI(3,4)P_2_ contributes to filopodia tip functions / organisation. Future work will aim at identifying the role of PI(3,4)P_2_ in filopodia as well as the proteins that are responsible for its accumulation at filopodia tips.

**Fig. 3.**
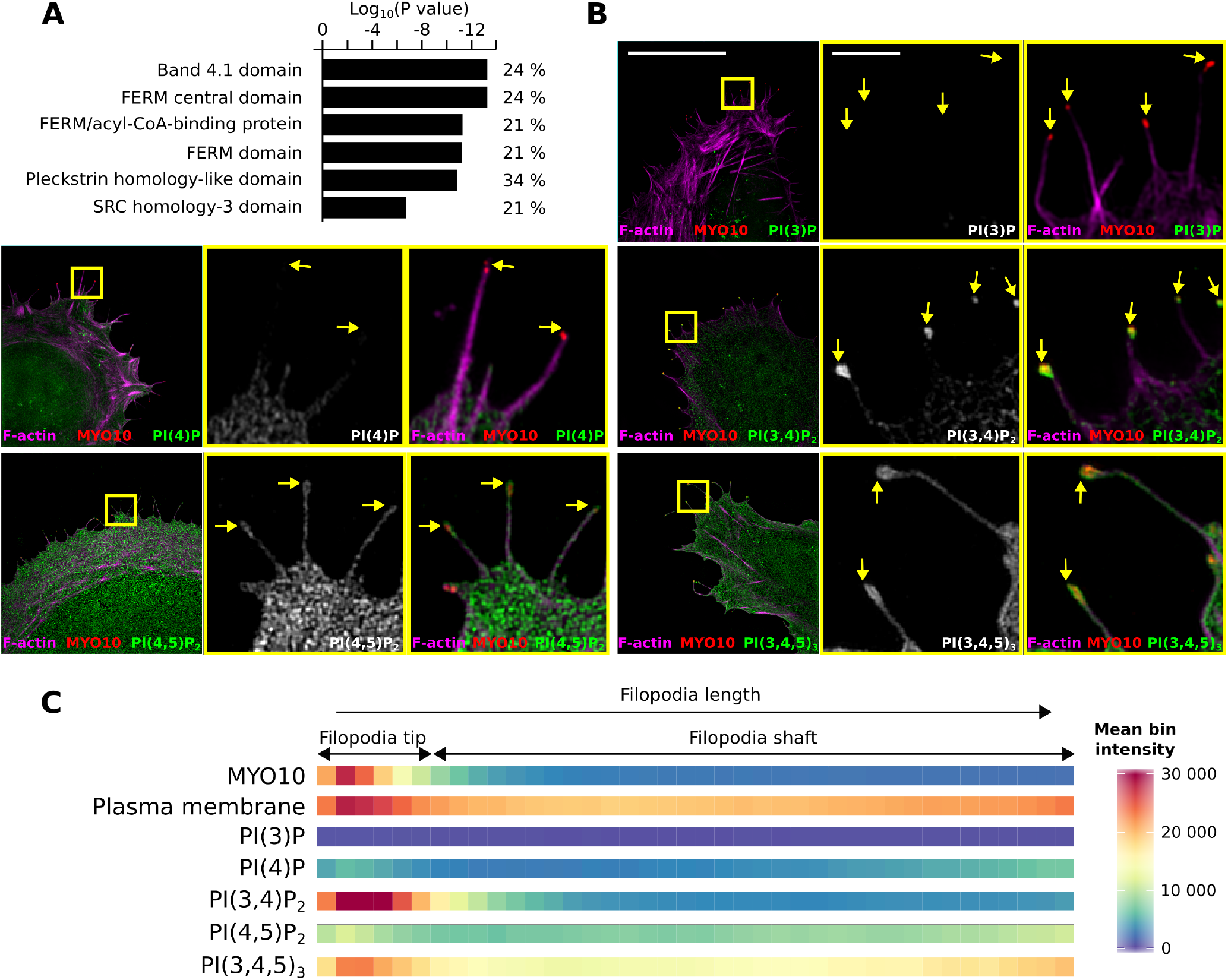
Mapping of phosphoinositides within filopodia. A: Functional annotation analysis looking for protein domain enrichments within the core filopodia proteins identified here (detected in at least 60% of filopodia). This analysis was performed using the INTERPRO database integrated within the DAVID platform (Huang et al., 2009). B: To map the distribution of the various phosphoinositides (PI) in filopodia, GFP-tagged probes binding with high affinity to a single PI species were transiently expressed together with MYO10-mScarlet in U2OS cells (PI species, probe used; PI(3)P, GFP-FYVE-PH; PI(4)P, GFP-P4M; PI(4,5)P_2_, GFP-PLC(Γ1)-PH; PI(3,4,5)P_3_, GFP-BTK-PH; PI(3,4)P_2_ GFP-TAAP-PH). Cells were then plated on fibronectin for 2 h, fixed, stained for F-actin and imaged using SIM. Maximal projections are displayed. Scale bars: (main) 20 μm; (inset) 2 μm. C: The distribution of each probe within filopodia, was then analysed and displayed as previously described (Figure 1 and Figure 2).

### SIM-mapping reveals novel filopodia tip proteins and subdomains within filopodia shafts

To quantitatively analyse the preferential recruitment of the core filopodia proteins, identified here, to filopodia tips or shafts, an enrichment ratio was calculated (intensity at filopodia tips / intensity at filopodia shafts) (Figure 4, see methods for details). As expected, proteins known to compose the tip complex such as MYO10 (Berg and Cheney, 2002), VASP (Rottner et al., 1999), DIAPH3 (Yang et al., 2007), FMNL3 (Harris et al., 2010) and RAPH1 (Krause et al., 2004) were strongly enriched to filopodia tips (Figure 4). Importantly, our analysis revealed that F-actin is low in the filopodium tip compared to the filopodium shaft (Figure 1B, 2 and 4). Conversely, the amount of plasma membrane (labelled using CAAX-GFP) was slightly enriched at filopodium tips compared to shafts (Figure 2 and 4). This is likely due to the formation of a bud at the end of filopodia. Therefore, the enrichment score of a POI was compared to the enrichment score obtained for the plasma membrane rather than the enrichment score of F-actin (Figure 4). Importantly, our mapping revealed several original filopodia tip proteins such as p130Cas (BCAR1), tensin-4 (TNS4) and adapter molecule crk (CRK). Other proteins displaying preferential recruitment to filopodia tips over filopodia shafts include the integrin activity modulators TLN1 and talin-2 (TLN2), kindlin-2 (FERMT2) and ICAP-1 (ITGB1BP1) (Figure 2 and 4). Proteins strongly enriched to filopodia shafts include predominantly actin regulating and cross-linking proteins such as eplins (LIMA1a and LIMA1b), alpha-actinin-1 (ACTN1), plastin (PLS3), and ezrin (EZR) (Figure 2 and 4). However, not all the actin regulators follow the F-actin distribution within filopodia. Indeed, some actin binding proteins such as cofilin (CFL1), FSCN1 or cortactin (CTTN) are more evenly distributed throughout the filopodia while others such as FMLN3, VASP or DIAPH3 are strongly enriched to the tips (Figure 2 and 4). Interestingly, epidermal growth factor receptor kinase substrate 8 (EPS8) and IRSp53 (BAIAP2), two regulators of filopodia formation, which are also known to interact with each other (Liu et al., 2010), appear to be spatially segregated within established filopodia as BAIAP2 accumulates at filopodia tips while EPS8 is enriched to filopodia shafts (Figure 2 and 4). Importantly, filopodia shafts can be segmented into subdomains depending on the degree of penetration of certain proteins through the filopodium (from the base of the shaft towards the tip). For instance, ACTN1 labels only the base (first 20%) of the shaft while proteins such as EZR or LIMA1 label the first 50-60% portion of the shaft (Figure 2). The segregation of the filopodia shaft into subdomains could indicate that these segments are also functionally different.

**Fig. 4.**
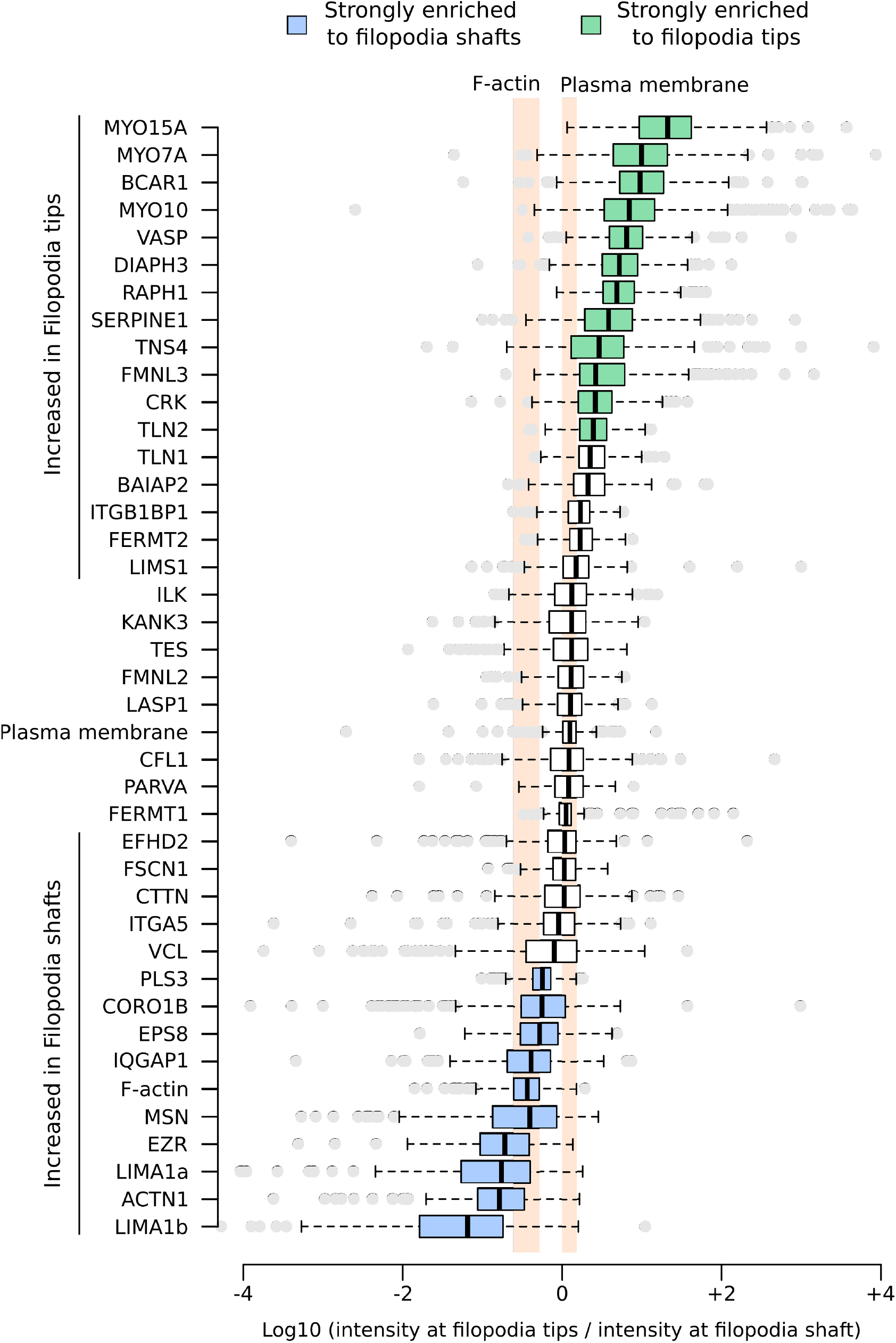
Preferential enrichment of adhesion proteins to filopodium shaft or tip. The preferential recruitment of the core filopodia proteins, identified here, to filopodia tips or shafts, was assessed by calculating an enrichment ratio (average intensity at filopodium tip / averaged intensity in the shaft). Results are displayed as Tukey box plots using a logarithmic scale and the POI are ordered as a function of the median of their enrichment score. The Tukey box plots represent the median and the 25th and 75th percentiles (interquartile range); points are displayed as outliers (represented by dots) if 1.5 times above or below the interquartile range (represented by whiskers). The enrichment scores of F-actin and of the plasma membrane (labelled by CAAX-GFP) are highlighted in orange. POI with a median enrichment score displaying at least a two-fold change over the median enrichment score of the plasma membrane were considered to be either strongly enriched to filopodia tips (highlighted in green) or strongly enriched to filopodia shafts (highlighted in blue). Proteins displaying enrichment scores statistically different from the enrichment scores of the plasma membrane are labelled as either “increased in filopodia tips” or “increased in filopodia shafts”.

### Filopodia adhesions are distinct from other adhesions

In migrating cells, adhesion sites are highly dynamic structures that undergo a well-defined force-dependent maturation sequence from nascent adhesions to fibrillar adhesions (Gardel et al., 2010). Our SIM based mapping allowed us to identify core filopodia proteins (detected in more than 60% of filopodia) including several established integrin binders and integrin activity regulators (talins, TLN1 and TLN2; kindlins, FERMT1 and FERMT2; ITGB1BP1), while other integrin binders and regulators such as tensin-1 (TNS1), tensin-3 (TNS3), MDGI (FABP3), filamin-a (FLNA) and nischarin (NISCH) were absent from filopodia. In addition, components of the signalling modules BCAR1-CRK and the Ilk-pinch-parvin complex (ILK, LIMS1 and PARVA; IPP complex) were detected in filopodia with similar distributions (Figure 1 and 2, Figure S1-S11). Several proteins accumulating in FA were also detected in most filopodia including VASP, ACTN1, testin (TES), LASP-1 (LASP1) and vinculin (VCL). Interestingly, within the same protein family, the ability of a protein to localise to filopodia can be isoform specific. For instance, only tensin-4 (TNS4) and KANK3 were found to localise to filopodia while the other family members did not (Figure 1 and 2, Figure S1-S11).

Filopodia are regions of low force in the cell (Bornschlögl, 2013) and therefore it is not surprising that proteins associated with mature and/or fibrillar adhesions were not identified as core filopodia proteins. These proteins include the actin-binding tensin isoforms (TNS1-3) (Zaidel-Bar et al., 2003), PDLIM1/5/7 (Kuo et al., 2011; Schiller et al., 2011), TRIP6 (Kuo et al., 2011; Schiller et al., 2011), Zyxin (ZYX) (Zaidel-Bar et al., 2003) and palladin (PALLD) (Azatov et al., 2016) (Figure 1 and 2, Figure S1-S11). In addition, the major non-muscle myosins mediating cellular contractility, non muscle myosin heavy chain IIa (MYH9) and non muscle myosin heavy chain IIb (MYH10), were not detected in filopodia. Importantly, multiple proteins associated with the low force-baring nascent adhesions including paxillin (PXN) (Pasapera et al., 2010), FAK (PTK2) (Lawson et al., 2012) or actin-related protein 3 (ACTR3, also known as ARP3) (Swaminathan et al., 2016), were classified as filopodia accessory, not core, proteins as they were detected in less than 40% of filopodia (Figure 1 and 2, Figure S1-S11). Thus, our SIM mapping indicates that filopodia adhesions consist of a unique set of proteins, the filopodome, and are distinct from classical nascent adhesions, focal adhesions and fibrillar adhesions.

The absence of PXN or PTK2 in more than 60% of filopodia was unexpected as there are documented interactions between these two proteins and a large number of the core filopodia proteins identified here (Figure S11B). Therefore, we validated the mapping by staining endogenous proteins in RFP-MYO10 expressing cells. In line with our mapping data from the GFP-tagged proteins, we found that endogenous ILK (Figure S12A) and TLN1 (Figure S12B) localise to filopodia. PXN (Figure 5A) and PTK2 (Figure S12C) were observed only in a small percentage of filopodia. When PXN was detected within filopodia, its localisation varied between being detected at the tip and / or in the shaft (Figure 5A).

**Fig. 5.**
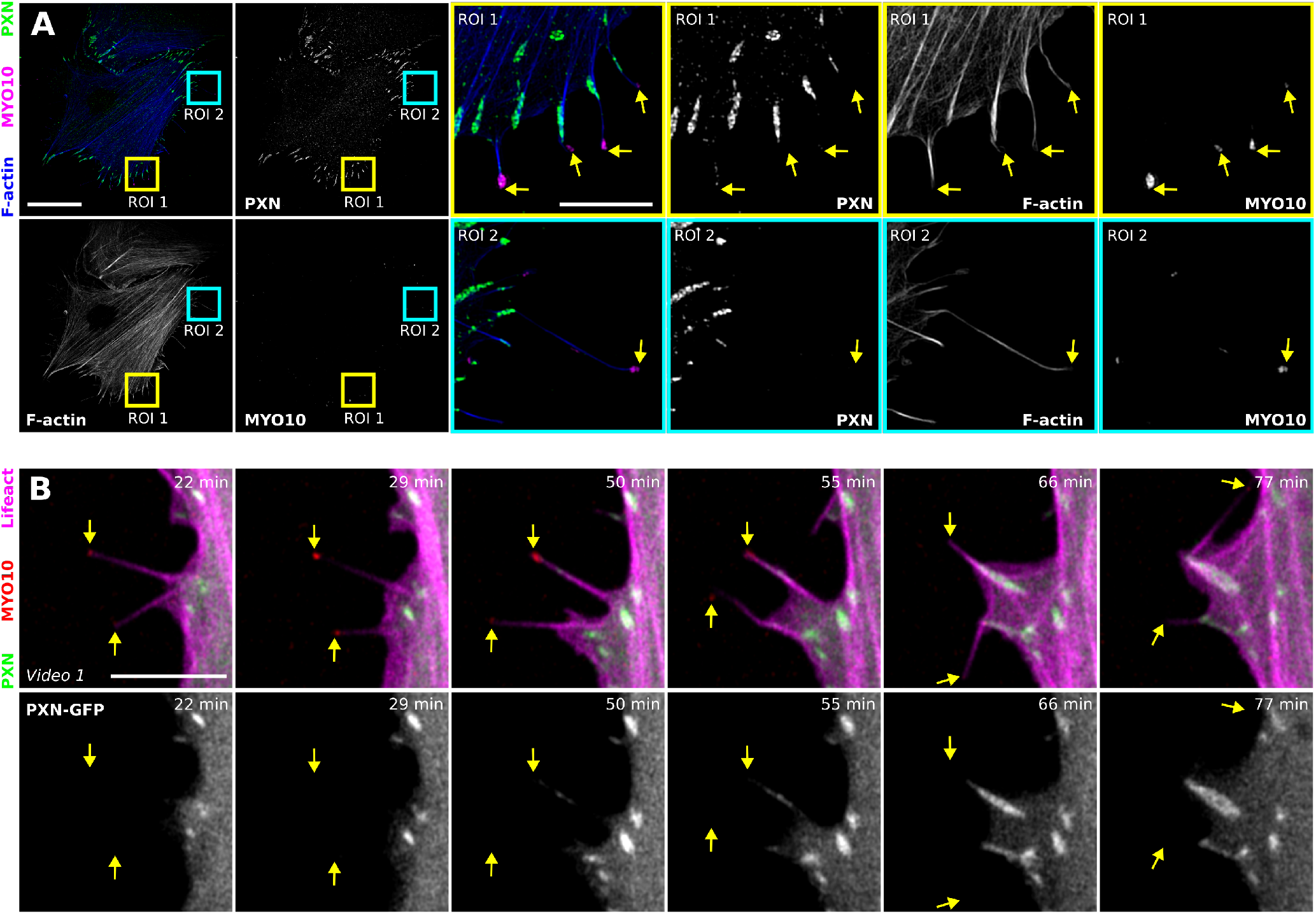
Filopodia tip adhesions nucleate nascent adhesions. A: U2OS cells expressing MYO10-mScarlet were plated on fibronectin for 2 h, fixed and stained for endogenous PXN and F-actin before being imaged using SIM. A representative maximum intensity projection is displayed. The blue and yellow squares highlight ROI, which are magnified. Scale bars: (main) 20 μm; (inset) 5 μm. B: U2OS cells transiently expressing lifeact-mTurquoise2, PAX-mEmerald and MYO10-mScarlet were plated on fibronectin and imaged live using an Airyscan confocal microscope (1 picture every 30 s; scale bar, 5 μm). The full video is available as supplementary information (Video 1). Images displayed highlight time points of interest in a magnified area. Yellow arrows highlight filopodia tips.

### Filopodia adhesions nucleate nascent adhesions

In fixed cells, individual filopodia represent the entire spectrum of filopodia dynamics from newly forming to stable and mature filopodia. We hypothesised that the variable localisation of PXN to filopodia could be linked to filopodia dynamics and therefore imaged PXN-GFP in U2OS cells co-expressing CFP-lifeact and RFP-MYO10 using an Airyscan confocal microscope (Figure 5B and Video 1). Live cell imaging revealed that unstable filopodia were devoid of PXN. Instead, PXN was recruited to filopodia following filopodia stabilisation, initially detected briefly at the tip, followed by localisation to, and increased clustering in, filopodia shafts, possibly due to actin retrograde flow in filopodia (Figure 5B and Video 1). Upon lamellipodia advancement, these clusters of PXN gave rise to FA (Figure 5B and Video 1). These data suggest that stabilized filopodia adhesions can trigger the nucleation of nascent adhesions. This is consistent with our previous work demonstrating that filopodia stabilization precedes FA maturation and the overall hypothesis that cycles of filopodia stabilization and FA maturation direct cell migration (Jacquemet et al., 2016).

### BCAR1 is a novel component of the filopodia tip complex

Talin-mediated integrin activation and linkage to the actin cytoskeleton are required for filopodia stabilization and maturation into adhesions. Adhesion maturation strongly correlates with increasing forces ranging from the modest forces of individual filopodia (5–25 pN with a maximum potential of up to 2 nN) (Bornschlögl, 2013; Albuschies and Vogel, 2013; Leijnse et al., 2015; Bornschlögl et al., 2013; Alieva et al., 2018) to the strong actomyosin-mediated contractile forces exerted at FA in 2D that are in the nano-Newton range (Galbraith et al., 2002). This prompted us to review our core filopodia proteins for potential force mediators. BCAR1 (p130Cas), one of our filopodia core components (Figure 1) strongly enriched in filopodia tips (Figure 4), has been implicated as a mechanosensitive protein that becomes phosphorylated by SRC in response to mechanical stretch (Sawada et al., 2006). We first validated our mapping data by staining endogenous BCAR1 and found a similarly strong localisation to filopodia tips as GFP-BCAR1 (Figure 6A). In addition, a phosphorylation-specific antibody indicated that endogenous BCAR1 is phosphorylated at filopodia tips (Figure 6B and Figure S12D) where it co-localises with MYO10 but not with PXN (Figure 6B and Figure S12D). Next, we analysed the dynamics of BCAR1 within filopodia in U2OS cells transiently expressing CFP-lifeact, BCAR1-GFP and RFP-MYO10 imaged using an Airyscan confocal microscope (Figure 6C and Video 2). Live cell imaging revealed that, unlike PXN, BCAR1 is always found at filopodia tips regardless of filopodia stability (Figure 6C and Video 2). Altogether, our data indicate that BCAR1 is a constitutive component of the filopodia tip complex (Figure 1 and 6) and its phosphorylation status suggests that it may play a role in ECM sensing in filopodia adhesion.

**Fig. 6.**
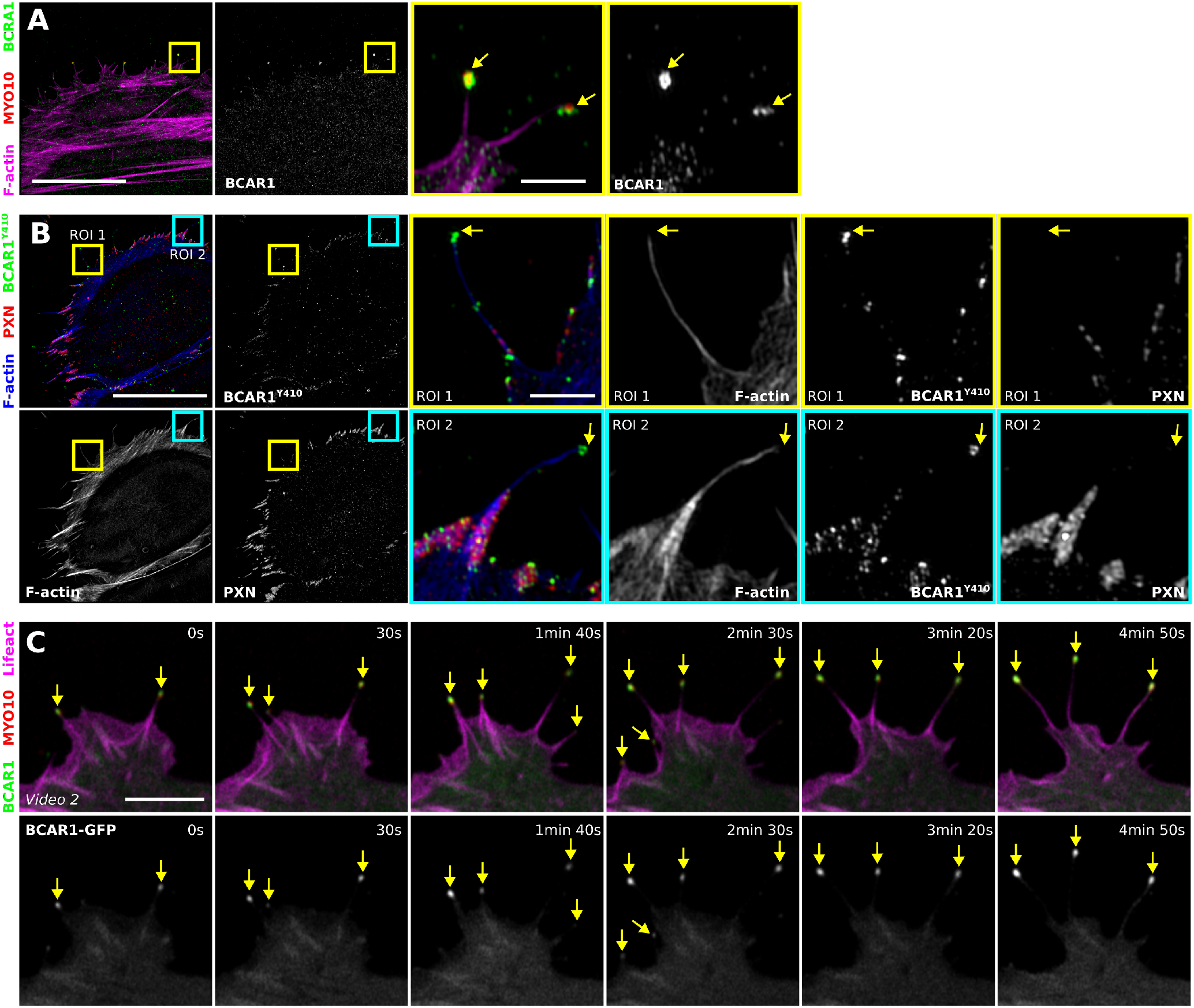
BCAR1 is part of the filopodia tip complex. A-B: U2OS cells expressing MYO10-mScarlet were plated on fibronectin for 2h, fixed and stained for F-actin and endogenous BCAR1 (A) or phospho-BCAR1 (Y410, B) before being imaged using SIM. Representative maximum intensity projections are displayed. The blue and yellow squares highlight ROI, which are magnified. Scale bars: (main) 20 μm; (inset) 2 μm. C: U2OS cells transiently expressing lifeact-mTurquoise2, BCAR1-eGFP and MYO10-mScarlet were plated on fibronectin and imaged live using an Airyscan confocal microscope (1 picture every 10 s; scale bar, 5 μm). The full video is available as supplementary information (Video 2). Images displayed highlight time points of interest in a magnified area. Yellow arrows highlight filopodia tips.

### BCAR1 is recruited to filopodia tips via its CCHD domain

BCAR1 localisation to FA has been suggested to be regulated via direct interaction with PXN or PTK2 (Zhang et al., 2017; Donato et al., 2010; Wang and McNiven, 2012), both of which are absent from most filopodia tips. In addition, BCAR1 recruitment to filopodia tips appears insensitive to inhibition of SRC, PTK2 or cellular contractility (Figure S13). Together, this suggested that BCAR1 localisation to filopodia tips occurs via a distinct mechanism to its FA targeting. To gain insight on how BCAR1 is recruited to filopodia tips, we sought to identify which domain of BCAR1 is required for filopodia targeting. BCAR1 is composed of five principal domains including an N-terminal SH3 domain followed by a proline-rich region, a substrate domain, a serine-rich region and a C-terminal Cas family homology domain (CCHD) (Janoštiak et al., 2014). Previous work demonstrated that both the SH3 and the CCHD domains were required to target BCAR1 efficiently to FA (Donato et al., 2010; Harte et al., 2000; Braniš et al., 2017). Therefore, we chose to map localisation of BCAR1 constructs lacking the SH3 and / or the CCHD domains within filopodia. As previously described, constructs lacking the SH3 or the CCHD domains accumulated poorly to FA while the construct lacking both domains was completely absent from FA (Figure 7A) (Harte et al., 2000; Braniš et al., 2017). Importantly, constructs lacking the CCHD domain failed to accumulate to filopodia tips while the construct lacking the SH3 domain accumulated to filopodia tips to the same extent as the full-length wild-type construct (Figure 7A and 7B). This indicates that BCAR1 CCHD domain, but not the SH3 domain, is required to target BCAR1 to filopodia tips.

**Fig. 7.**
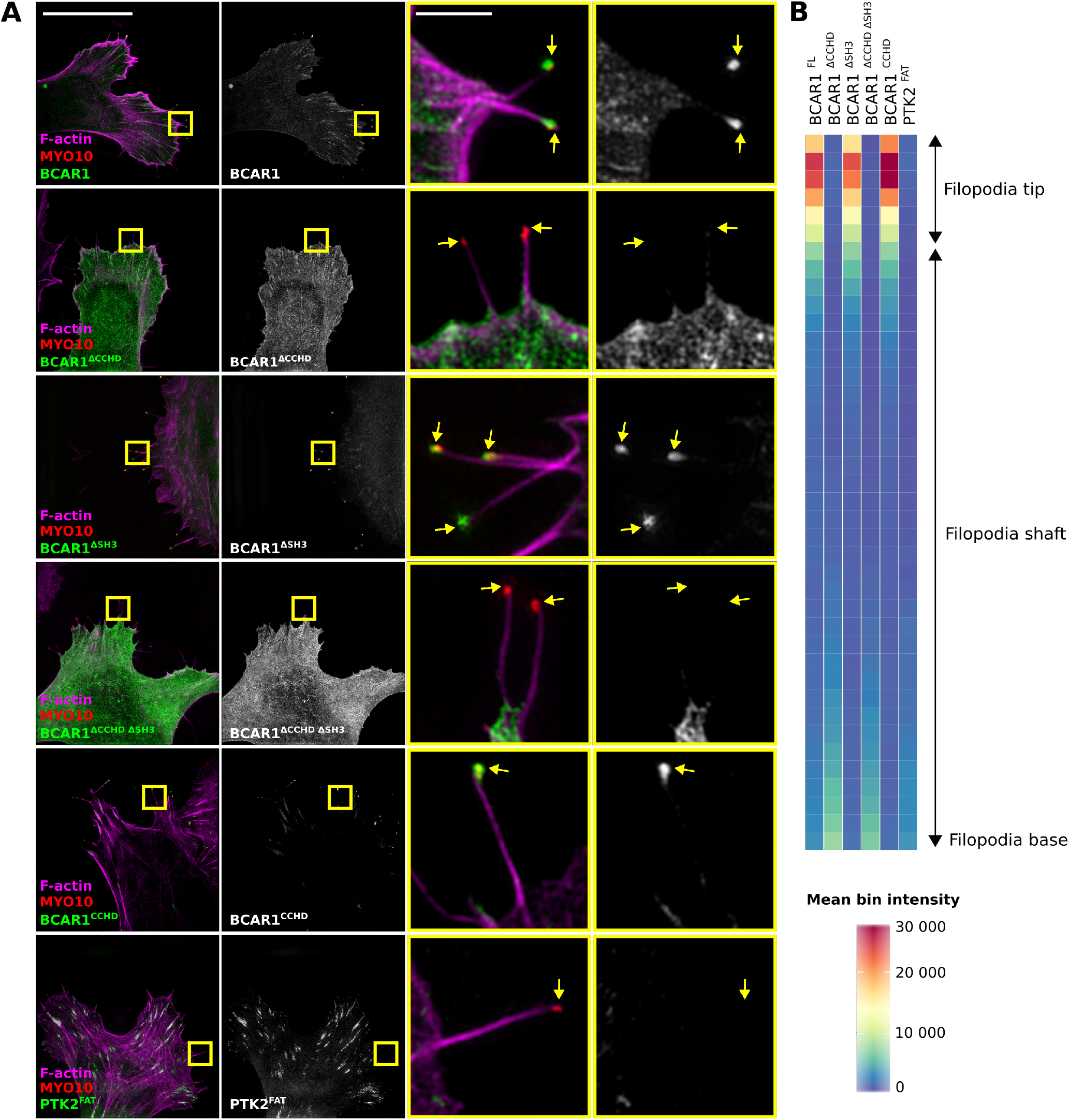
BCAR1 is recruited to filopodia tips via its CCHD domain. A: U2OS cells expressing MYO10-mScarlet together with GFP tagged-BCAR1 or various BCAR1 deletion constructs, or GFP-PTK2-FAT (PTK2^FAT^) were plated on fibronectin for 2 h, fixed, stained for F-actin and imaged using SIM. BCAR1 deletion constructs include a construct where BCAR1 SH3 domain was deleted (BCAR1^ΔSH3^), one construct where BCAR1 CCHD domain was deleted (BCAR1^ΔCCHD^), one construct where both BCAR1 SH3 and CCHD domains were deleted (BCAR1^ΔCCHDΔSH3^) and one construct where BCAR1 CCHD domain alone was expressed (BCAR1^CCHD^). For each condition, maximal intensity projections are displayed and yellow squares highlight ROI, which are magnified. Yellow arrows highlight filopodia tips. Scale bars: (main) 20 μm; (inset) 2 μm. B: The distribution of each construct within filopodia, was then analysed and displayed as previously described (Figure 1 and Figure 2).

We next sough to test if the BCAR1 CCHD domain was sufficient for filopodia tip localisation. BCAR1 CCHD domain is structurally very similar to the focal adhesion targeting (FAT) domain of PTK2 (Mace et al., 2011; Zhang et al., 2017). Therefore, the cellular distribution of the PTK2 FAT domain and BCAR1 CCHD was compared using SIM (Figure 7A). As expected, both BCAR1 CCHD and PTK2 FAT constructs accumulated in FA (Figure 7A). Strikingly, only BCAR1 CCHD, and not PTK2 FAT, accumulated to filopodia tips (Figure 7A and 7B). This demonstrates that BCAR1 CCHD is both a focal adhesion targeting domain and a filopodia tip complex targeting (FTCT) domain. To our knowledge, such a FTCT domain is an original finding and future work will aim at identifying the mechanism controlling BCAR1 accumulation to filopodia tips.

### BCAR1 contributes to filopodia stabilisation and stiffness sensing

Previously we identified that integrin activation as well as integrin downstream signalling are required for efficient filopodia formation and stabilisation (Jacquemet et al., 2016). In particular silencing of TNL1 or inhibition of SRC strongly inhibit filopodia formation (Jacquemet et al., 2016). In order to investigate the potential role of BCAR1 in this process, we silenced BCAR1 expression using two independent siRNAs (Figure 8A) in both U2OS and MDA-MB-231 cells transiently expressing MYO10-GFP (Figure 8B and 8C). In both cell types, silencing of BCAR1 led to a small increase in the number filopodia, indicating that BCAR1 is not essential for filopodia but may modulate their formation (Figure 8B and 8C). We next assessed the role of BCAR1 in regulating filopodia dynamics. Strikingly, BCAR1-silenced cells displayed a higher proportion of unstable filopodia and a lower proportion of stable filopodia compared with control cells, indicating that BCAR1 contributes to filopodia stabilisation (Figure 8D).

**Fig. 8.**
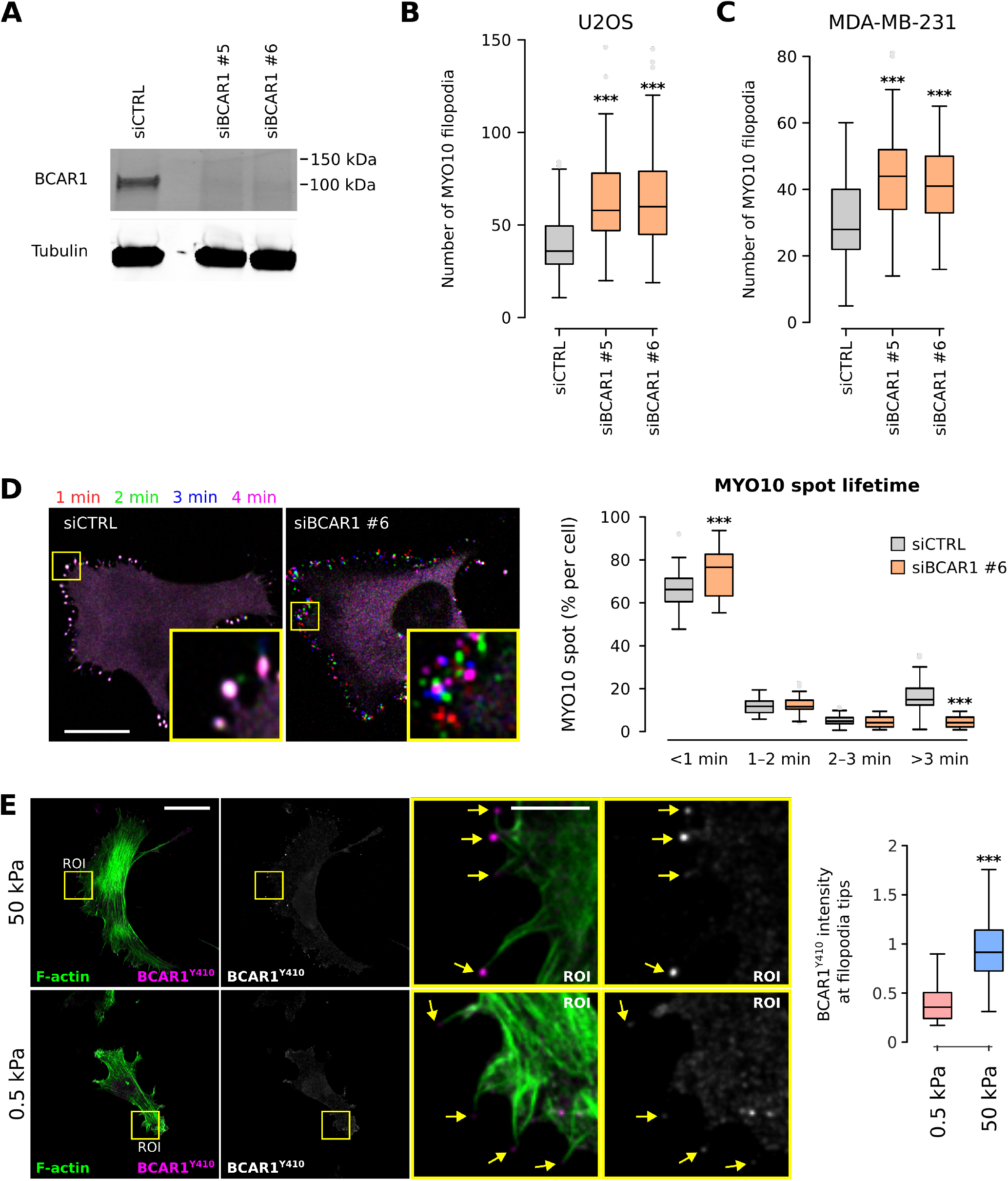
BCAR1 regulate environment sensing at filopodia tips. A: U2OS cells previously silenced for BCAR1 (oligo #5 and #6) were lysed and BCAR1 protein levels were assessed by western blot. B-C: U2OS (B) and MDA-MB-231 (C) cells previously silenced for BCAR1, using two distinct siRNAs, and transiently expressing MYO10-GFP, were plated on fibronectin for 2 h, fixed and the number of MYO10-positive filopodia per cell was quantified (n > 65 cells, three biological repeats; ***P value ≤ 5.4 x10^−6^). D: U2OS cells previously silenced for BCAR1 (oligo #6) and transiently expressing MYO10-eGFP were plated on fibronectin and imaged live using an Airyscan confocal microscope (1 picture every 5 s; scale bar, 20 μm). Representative images at different time points are shown. For each condition, MYO10-positive particles were automatically tracked and MYO10 spot lifetime (calculated as a percentage of the total number of filopodia generated per cell) was plotted and displayed as Tukey box plots (see method for details; three biological repeats, more than 40 cells per condition, ***P value ≤ 8.78 x10^−5^). E: U2OS cells expressing MYO10-mScarlet were plated on fibronectin-coated polyacrylamide gels of defined stiffness (0.5 kPa, soft; 50 kPa, stiff) for 2 h. After fixation, cells were stained for F-actin and phospho-BCAR1 (Y410) before being imaged using a spinning disk confocal microscope. Maximal intensity projections are displayed and yellow squares highlight ROI, which are magnified. Yellow arrows highlight filopodia tips. Scale bars: (main) 20 μm; (inset) 5 μm.

BCAR1 phosphorylation triggers the recruitment of multiple proteins required for activation of signalling pathways driving cell migration (Janoštiak et al., 2014). BCAR1 Y410 phosphorylation is increased upon mechanical stretch, et al., 2007) EPS8 (Calderwood et al., 2003) and ACTN1 (Tadokoro et al., 2011), were found to localise to filopodia. Therefore, future challenges will be to establish how integrin functions including transport and activation are coordinated by multiple adaptors within filopodia, and what is the sequence and hierarchy of binding of these proteins to their overlapping binding sites on the short cytoplasmic domains of integrin β-subunits.

Initial integrin-ECM engagement leads to the activation of several signalling nexus including SRC, FAK (PTK2), and ILK. Here we identified that the ILK-pinch (LIMS1)-parvin (PARVA) (IPP) complex localises to filopodia (Figure 1 and 2). ILK associates with FERMT2 and therefore FERMT2 (Montanez et al., 2008) could recruit ILK and the IPP complex to filopodia. The contribution of ILK, LIMS1 or PARVA to filopodia function are not known but previous work reported a role for ILK and PARVA in the formation of filopodium-like protrusions in 3D (Shibue et al., 2013). In this context, both ILK and PARVA contributed to cancer cell metastatic outgrowth and survival at distal sites (Shibue et al., 2013). SRC kinase is a major regulator of filopodia formation and stabilisation and active SRC localises to filopodia (Jacquemet et al., 2016). At FA, principal SRC substrates include PTK2, paxillin (PXN) and p130Cas (BCAR1). Here we report that PTK2 and PXN are absent from initial filopodia adhesions (Figure 1, 2 and 5). However, BCAR1 strongly accumulates very early on in filopodia at tips of newly forming filopodia structures (Figure 1, 2, 4, 6, 7 and 8). BCAR1 is not an essential regulator of filopodia formation but rather regulates filopodia stabilization (Figure 8). The recruitment of BCAR1 to filopodia tips is mediated by the CCHD domain but the distinction between targeting of CCHD to filopodia adhesions and FA remains to be investigated. Overall, BCAR1 localisation to filopodia and FA appears to be regulated by different molecular interactions. While PTK2 and PXN (absent from newly formed filopodia) and the BCAR1 SH3 domain are required for BCAR1 FA targeting (Donato et al., 2010; Wang and McNiven, 2012; Zhang et al., 2017), they are dispensable for BCAR1 recruitment to filopodia tips.

Filopodia sense ECM topography and/or ECM stiffness (Albuschies and Vogel, 2013; Chan and Odde, 2008; Wong et al., 2014). We previously observed that filopodia stabilization precedes FA maturation and that this process directs cell migration (Jacquemet et al., 2016). Here, we observed that PXN-positive adhesions can form in filopodia shafts which, upon lamellipodia advancement (Hu et al., 2014), leads to the formation of FA (Figure 5). As filopodia are widely used by cells migrating in fibrillar matrices (Jacquemet et al., 2013; Paul et al., 2015), it is tempting to speculate that the formation of PXN-positive adhesions in the filopodia shaft could be a mechanism by which cells sense matrix alignment. In the context of stiffness sensing, filopodia tip proteins TLN1 and BCAR1 have both previously been reported to be mechanosensitive (Janoštiak et al., 2014). Forces exerted by individual filopodia typically range from 5–25 pN (with a maximum potential of up to 2 nN) and are principally mediated by actin retrograde flow (Bornschlögl, 2013; Bornschlögl et al., 2013; Leijnse et al., 2015) and, upon maturation, by myosin contractility (Alieva et al., 2018). Accordingly, neither MYH9 or MYH10 were detected within filopodia (Figure 1 and 2) but rather these proteins can cluster at the base of filopodia (Alieva et al., 2018). In the case of TLN1, forces above 5 pN (15 pN for further activation) can induce conformational changes that trigger a switch from talin–RIAM to talin–vinculin complexes leading to FA stabilisation (Yao et al., 2016). In the case of BCAR1, forces induce the stretching of BCAR1 substrate domain leading to its phosphorylation by SRC (Sawada et al., 2006). BCAR1 phosphorylation is not mediated by actomyosin contractility but instead depends on an intact actin cytoskeleton (Hotta et al., 2014). Importantly, here we demonstrate that BCAR1 phosphorylation at filopodia tips is stiffness sensitive (Figure 8D). While the exact amount of force required to activate BCAR1 remains unknown, BCAR1 substrate domain is disorganised and it is likely that the weak forces mediated by actin retrograde flow alone (5 pN in magnitude) are sufficient to mediate activation (Hotta et al., 2014). Due to the low force requirement for BCAR1 activation compared to TLN1, it is likely that, at the filopodia tip, BCAR1 will be one of the first mechanosensitive proteins to be activated. BCAR1 activation may then lead to activation of Rap1 (Kirsch et al., 1998; Gotoh et al., 2000), which would in turn promote talin-mediated integrin activation, adhesion re-enforcement and filopodia stabilisation.

Altogether, we have revealed that filopodia adhesions consist of a unique set of proteins, the filopodome, that are distinct from classical nascent adhesions, FA and fibrillar adhesions. Our mapping will be a valuable resource for future studies aimed at unravelling the biological relevance of filopodia in different developmental and pathological conditions involving cell motility and cellular responsiveness to environmental cues.

## Materials and Methods

### Cell culture and transient transfection

U2OS osteosarcoma cells and MDA-MB-231 (triple-negative human breast adenocarcinoma) cells were grown in DMEM supplemented with 10% FCS. U2OS cells were purchased from DSMZ (Leibniz Institute DSMZ-German Collection of Microorganisms and Cell Cultures, Braunschweig DE, ACC 785). MDA-MB-231 cells were provided by ATCC. All cells were tested for mycoplasma contamination. Plasmids of interest were transfected using Lipofectamine 3000 and the P3000TM Enhancer Reagent (Thermo Fisher Scientific) according to the manufacturer’s instructions. The expression of proteins of interest was suppressed using 100 nM siRNA and lipofectamine 3000 (ThermoFisher Scientific) according to manufacturer’s instructions. The siRNA used as control (siCTRL) was Allstars negative control siRNA (Qiagen, Cat. No. 1027281). The siRNAs targeting BCAR1 were purchased from Qiagen (siBCAR1#5, Hs_BCAR1_5 FlexiTube siRNA, Cat. No. SI02757734; siBCAR1#6, Hs_BCAR1_6 FlexiTube siRNA, Cat. No. SI02757741).

### Reagents, antibodies and compounds

Mouse monoclonal antibodies used in this study were against p130Cas (BCAR1, Santa Cruz Biotechnology, SC-20029; 1:100 for immunofluorescence (IF), 1:1000 for western blotting (WB)), talin-1 (TLN1, Sigma, clone 8d4, Cat. No. T3287; 1:100 for IF), FAK (PTK2, BD Biosciences, Clone 77, Cat. No. 610087; 1:100 for IF), α-tubulin (Hybridoma Bank, clone 12G10; 1:1000 for WB) and paxillin (PXN, BD Biosciences, Clone 349, Cat. No. 610051; 1:100 for IF). Rabbit polyclonal antibody raised against Phosphop130Cas (BCAR1, Tyr410) was purchased from Cell signalling (Cat. No. 4011). Rabbit monoclonal antibody raised against human ILK (EPR1592, Cat. No. ab76468; 1:100 for IF) was purchased from Abcam. Alexa Fluor 488, Alexa Fluor 568 phalloidin and Atto 647 phalloidin were provided by Thermo Fisher Scientific. SiR-actin was provided by Cytoskeleton (Cat. No. CY-SC001). Bovine plasma fibronectin was provided by Merck (341631). Dimethyl-sulphoxide (DMSO; D2650) was obtained from Sigma. The SRC (PP2, S7008) and FAK (PTK2) inhibitors (PF-573228, S2013) were provided by Selleckchem. The myosin II inhibitor (blebbistatin, 72402) was purchased from STEM-CELL Technologies.

### Plasmids

The following plasmids were described previously eGFP-TNS1, eGFP-TNS2, eGFP-TNS3 (Georgiadou et al., 2017), eGFP-TNS4 (Muharram et al., 2014), GFP-MDGI (FABP3) (Nevo et al., 2009) and GFP-Sharpin (SHARPIN) (Rantala et al., 2011). GFP-FAK-FAT and GFP-FL-FAK were a gift from David D. Schlaepfer (UC San Diego Health, US). EFHD2-GFP was a gift from Dirk Mielenz (University of Erlangen-Nuremberg, DE) (Avramidou et al., 2007). CAAX-GFP was a gift from Gregory Giannone (Bordeaux University, FR). PEGFP-Talin-1 (TLN1), mCherry-Talin-2 (TLN2) and pDsRedC1-Kindlin-1 (FERMT1) were gifts from Ben Goult (University of Kent, UK). FMNL2-GFP was a gift from Robert Grosse (University of Marburg, DE) (Grikscheit et al., 2015). FMNL3-GFP (Harris et al., 2010) was a gift from Henry Higgs (Geisel School of Medicine at Dartmouth, US). PPFIA1-GFP was a gift from Guido Serini (University of Torino, IT). pEGFP-C1-Lamellipodin (RAPH1) was a gift from Matthias Krause (King’s College London, UK) (Krause et al., 2004). pEGFP-C2-Myo15a (MYO15A) was a gift from Jonathan Bird (NIH, Bethesda US) (Belyantseva et al., 2005). GFP-ICAP-1 (ITGB1BP1) was a gift from Daniel Bouvard (University of Grenoble, FR). GFP-KANK1, GFP-KANK2, GFP-KANK3 and GFP-KANK4 plasmids were gifts from Reinhard Fässler (Max Planck Institute of Biochemistry, Martinsried, DE) (Sun et al., 2016). Plasmids encoding BTK-PH-EGFP and PLC(δ1)-PH-EGFP were gifts from Matthias Wymann (University of Basel, Switzerland). The EGFP-tagged tandem FYVE was a gift from Harald Stenmark (Oslo University Hospital) (Gillooly et al., 2000). Plasmids encoding BCAR1 deletion constructs (GFP-Cas-wt, GFP-CasdeltaCCH, GFP-CasdeltaSH3 and GFP-Cas-deltaCCH-deltaSH3) were gifts from Daniel Rösel (Charles University in Prague, Czech Republic) (Braniš et al., 2017). Plasmids encoding Kindlin-2-GFP (FERMT2), Ezrin-GFP (EZR) and Vinculin-GFP (VCL) were gifts from Maddy Parsons (King’s College London, UK). The Moesin-GFP (MSN) plasmid was a gift from Buzz Baum (University College London, UK). Lifeact-mTurquoise2 was a gift from Joachim Goedhart (University of Amsterdam, NL) (Chertkova et al., 2017). Integrin alpha5-GFP (ITGA5) was a gift from Rick Horwitz (Allen institute for cell science, US).

The following plasmids were provided by Addgene (gift from, Addgene plasmid number): mEmerald-Alpha-Actinin-19 (Michael Davidson, 53989), mEmerald-Fascin-C-10 (Michael Davidson, 54094), pGFP Cas (Kenneth Yamada, 50729), mEmerald-Cofilin-C-10 (Michael Davidson, 54047), mEmerald-Coronin1B-C-10 (Michael Davidson, 54049), pGFP CrkII (Kenneth Yamada, 50730), mEmerald-Cortactin-C-12 (Michael Davidson, 54051), mEmerald-mDia1-C-14 (Michael Davidson, 54156), mEmerald-mDia2-C-14 (Michael Davidson, 54158), mEmerald-Migfilin-C-14 (Michael Davidson, 54181), pEGFP-IQGAP1 (David Sacks, 30112) (Ren et al., 2005), mEmerald-LASP1-C-10 (Michael Davidson, 54141), EGFP-EPLIN alpha (Elizabeth Luna, 40947), EGFP-EPLIN beta (Elizabeth Luna, 40948), mEmerald-PINCH-C-14 (Michael Davidson, 54229), mEmerald-MyosinIIA-C-18 (Michael Davidson, 54190) (Burnette et al., 2014), mEmerald-MyosinIIB-C-18 (Michael Davidson, 54192), mEmerald-Palladin-C-7 (Michael Davidson, 54213), mEmerald-Parvin-C-14 (Michael Davidson, 54214), GFP-PTEN (Alonzo Ross, 13039) (Liu et al., 2005), mEmerald-Paxillin-22 (Michael Davidson, 54219) (Paszek et al., 2012), mEmerald-TESC-14 (Michael Davidson, 54276), mEmerald-VASP-N-10 (Michael Davidson, 54297), mEmerald-Zyxin-6 (Michael Davidson, 54319), E-cadherin-GFP (Jennifer Stow, 28009) (Miranda et al., 2001), pEGFP C1-Eps8 WT (Giorgio Scita, 74950) (Hertzog et al., 2010), GFP-P4M-SidM (Tamas Balla, 51469) (Hammond et al., 2014), pcDNA3.1-6His-MyoX (Emanuel Strehler, 47607) (Bennett and Strehler, 2008) and ARP3-GFP (Matthew Welch, 8462) (Welch et al., 1997).

The mScarlet-MYO10 construct was generated by inserting a gene block containing the mScarlet sequence (IDT, see supplements for sequence) (Bindels et al., 2017) into pcDNA3.1-6His-MyoX using the KpnI restriction site. The Cas CCHD construct was generated by inserting a gene block containing the BCAR1 CCHD sequence (IDT, see supplements for sequence) into pEGFP-C1 using the XhoI and BamHI restriction sites. Several of the GFP tagged constructs, used here, were generated by the Genome Biology Unit core facility cloning service (Research Programs Unit, HiLIFE Helsinki Institute of Life Science, Faculty of Medicine, University of Helsinki, Biocenter Finland) by transferring entry clones from the ORFeome collaboration library into mEmerald destination vectors using the standard LR reaction protocol. Entry clone (I.M.A.G.E. Consortium CloneID, (Lennon et al., 1996)) transferred into pcDNA6.2/N-emGFP-DEST include MYO7A (100069043), BAIAP2 (100006086), PDLIM5 (100003316), FERMT2 (100067023), FHL2 (100006502), FHL3 (100003225), PDLIM7 (100004765), PLS3 (100003800), TRIP6 (100005523), ALYREF (100066457), ANXA1 (100003803), BRIX1 (100004738), DIMT1 (100004112), FAU (100005039), HP1BP3 (100067555), PCOLCE (100004553), POLDIP3 (100000169), SERPINE1 (100003324) and SORBS1 (100064166). Entry clones transferred into pcDNA6.2/C-emGFP-DEST include TGM2 (100074142), P4HB (100073368), PDLIM1 (100069996), SYNCRIP (100074128), NISCH (100068156) and PRSS23 (100002084).

### SDS–PAGE and quantitative western blotting

Protein extracts were separated under denaturing conditions by SDS–PAGE and transferred to nitrocellulose membranes. Membranes were blocked for 1 h at room temperature with blocking buffer (LI-COR Biosciences) and then incubated overnight at 4°C with the appropriate primary antibody diluted in blocking buffer. Membranes were washed with PBS and then incubated with the appropriate fluorophore-conjugated secondary antibody diluted 1:5,000 in blocking buffer for 30 min. Membranes were washed in the dark and then scanned using an Odyssey infrared imaging system (LI-COR Biosciences). Band intensity was determined by digital densitometric analysis using the Odyssey software.

### Sample preparation for light microscopy

If not indicated otherwise, cells were plated on high tolerance glass-bottom dishes (MatTek Corporation, coverslip #1.7) pre-coated first with Poly-L lysine (10 μg/ml, 1 h at 37°C) and with bovine plasma fibronectin (10 μg/ml, 2 h at 37°C). To generate the filopodia map, U2OS cells transiently expressing a GFP-tagged protein of interest (POI) and MYO10-mScarlet were plated for 2 h on fibronectin-coated glass-bottom dishes. Samples were fixed and permeabilised simultaneously using a solution of 4% (wt/vol) PFA and 0.25% (vol/vol) Triton X-100 for 10 min. Cells were then washed with PBS, quenched using a solution of 1 M glycine for 30 min, and incubated with SiR-actin (100 nM in PBS) at 4°C until imaging (minimum length of staining, overnight at 4°C; maximum length, 1 week). Just before imaging, samples were washed three times in PBS and mounted in vectashield (Vectorlabs). To stain endogenous proteins, U2OS cells transiently expressing MYO10-mScarlet were plated on fibronectin-coated glass-bottom dishes. Samples were fixed and permeabilised simultaneously using a solution of 4% (wt/vol) PFA and 0.25% (vol/vol) Triton X-100 for 10 min. Cells were then washed with PBS, quenched using a solution of 1 M glycine for 30 min, and incubated with the primary antibody for 1 h (1:100). After three washes, cells were incubated with a secondary antibody for 1 h (1:100). Samples were then washes three times and stored in PBS or in PBS containing an actin stain (as indicated) at 4°C until imaging. Just before imaging, samples were washed three times in PBS and mounted in vectashield (Vectorlabs). All live-cell imaging experiments were performed in normal growth media, supplemented with 50 mM Hepes, at 37°C and in the presence of 5% CO_2_.

### Microscopy setup

The structured illumination microscope (SIM) used was DeltaVision OMX v4 (GE Healthcare Life Sciences) fitted with a 60× Plan-Apochromat objective lens, 1.42 NA (immersion oil RI of 1.516) used in SIM illumination mode (five phases × three rotations). Emitted light was collected on a front illuminated pco.edge sCMOS (pixel size 6.5 μm, readout speed 95 MHz; PCO AG) controlled by SoftWorx. The spinning disk microscope used was a Marianas spinning disk imaging system with a Yokogawa CSU-1 scanning unit on an inverted Zeiss Axio Observer Z1 microscope controlled by SlideBook 6 (Intelligent Imaging Innovations, Inc.). Objectives used were a long-working-distance 63× water (NA 1.15 water, LDC-Apochromat, M27) objective or a 100× (NA 1.4 oil, Plan-Apochromat, M27) objective. Images were acquired using either an Orca Flash4 sCMOS camera (chip size 2,048 x 2,048; Binning 2×2 enabled; Hamamatsu Photonics) or an Evolve 512 EMCCD camera (chip size 512 × 512; Photometrics). The confocal microscope used was a laser scanning confocal microscope LSM880 (Zeiss) with a 40x (1.4 oil) equipped with an Airyscan detector (Carl Zeiss). The microscope was controlled using Zen Black (2.3) and the Airyscan was used in standard super-resolution mode.

### Filopodia formation assays on fibronectin

Cells expressing human MYO10-GFP (to visualise filopodia tips) were plated for 2 h in full medium on glass-bottom dishes (MatTek Corporation) precoated with fibronectin (10 μg/ml). Cells were then fixed using PFA, washed with PBS, permeabilized and stained using phalloidin. Images were acquired on an SDC microscope using a 100x objective and the number of filopodia per cell was manually scored using Fiji (Schindelin et al., 2012; Rueden et al., 2017).

### Filopodia stability assay

To study the role of BCAR1 in filopodia stability, U2OS cells expressing MYO10-GFP were plated for at least 2 h on fibronectin before the start of live imaging (pictures taken every 5 s at 37oC, on an Airyscan microscope using a 40x objective). Filopodia lifetimes were then measured by identifying and tracking all MYO10 spots using the Fiji plugin TrackMate (Tinevez et al., 2017). In Trackmate, the LoG detector (estimated bob diameter = 0.8 μm; threshold = 20; subpixel localization enabled) and the simple LAP tracker (linking max distance = 1 μm; gap-closing max distance = 1 μm; gap-closing max frame gap = 0) were used.

### Mapping of proteins within filopodia

To map the localisation of each POI within filopodia, the images were first processed in Fiji (Schindelin et al., 2012) and the data analysed using R. Briefly, in Fiji, the brightness/contrast of each image was automatically adjusted using, as an upper maximum, the brightest cellular structure labelled in the field of view. In Fiji, line intensity profiles were manually drawn from filopodium tip to base and exported for further analysis. The Fiji script used to process the data is available as supplementary information (Script 1). For each POI, line intensity profiles were then compiled and analysed in R. To homogenise filopodia length, each line intensity profile was binned into 40 bins (using the median value of the pixels comprised in each bin). Using the line intensity profiles, the percentage of filopodia positive for each POI was quantified. A positive identification was defined as requiring at least three bins (out of 40), each with a minimum value of 10000 (bin values between 0-65535). The map of each POI was created by averaging hundreds of binned intensity profiles. The R script used to bin and average line intensity profiles is available as supplementary information (Script 2). The averaged binned intensity profiles of each POI is available in Table S2. The filopodia maps were then displayed as heatmaps in R (Script 3).

### Statistical analysis

The Tukey box plots represent the median and the 25th and 75th percentiles (interquartile range); points are displayed as outliers (represented by dots) if 1.5 times above or below the interquartile range (represented by whiskers). Box plots were generated using the online tool BoxPlotR (http://shiny.chemgrid.org/boxplotr/). Statistical analyses were performed when appropriate, and p-values are indicated in the figure legends. Unless otherwise indicated, the Student’s t test was used (unpaired, two tailed, and unequal variance, performed within LibreOffice Calc).

## Data availability

The authors declare that the data supporting the findings of this study are available within the article and from the authors on request.

## Supplemental material

**Figure S1 to Figure S11** show representative images highlighting the localisation of each of the proteins of interest (POI, arranged alphabetically) imaged to generate the filopodia map displayed in figure 2. **Figure S12** shows representative images of the endogenous staining of TNL1/2, PTK2, ILK and BCAR1 pY410. **Figure S13** shows that BCAR1 recruitment to filopodia tips is not affected by inhibition of SRC, FAK or myosin-II. **Movie 1** highlights PXN dynamics within filopodia during cell migration. **Movie 2** highlights BCAR1 dynamics within filopodia. **Table S1** highlights the various proteins / constructs imaged in this study as well as the number of filopodia analysed to generate Figures 1, 2, 3, 4 and 7. **Table S2** contains the numerical values used to generate the filopodia map displayed in Figure 2. **Script 1** is the ImageJ macro used to measure and export the line intensity profiles from filopodia. **Script 2** is the R code used to extract, compile and average the line intensity profiles previously measured in ImageJ. **Script 3** is the R code used to generate the filopodia map (using Table S2 as input) displayed in Figure 2.

## Acknowledgements

We thank J. Siivonen and P. Laasola for technical assistance and M. Saari for help with the microscopes. We thank Dirk Mielenz, Gregory Giannone, Ben Goult, Robert Grosse, Henry Higgs, Guido Serini, Matthias Krause, Jonathan Bird, Daniel Bouvard, Reinhard Fässler, Matthias Wymann, Harald Stenmark, Daniel Rösel, Maddy Parsons, Buzz Baum, Joachim Goedhart, Rick Horwitz, Michael Davidson, Kenneth Yamada, David Sacks, Elizabeth Luna, Alonzo Ross, Jennifer Stow, Giorgio Scita, Tamas Balla, Emanuel Strehler and Matthew Welch for providing reagents.

The Cell Imaging Core (Turku Centre for Biotechnology, University of Turku, Åbo Akademi University and Biocenter Finland) and the Genome Biology Unit (Research Programs Unit, HiLIFE Helsinki Institute of Life Science, Faculty of Medicine, University of Helsinki, Biocenter Finland) are acknowledged for services, instrumentation, and expertise.

This study has been supported by the Academy of Finland (G.J. and J.I.), Academy of Finland CoE for Translational Cancer Research (J.I.), ERC CoG grant 615258, Sigrid Juselius Foundation and the Finnish Cancer Organization (J.I.). M.M. has been supported by the Turku Drug Research Doctoral Programme.

## Authors contributions

Conceptualization, G.J. and J.I.; Methodology, G.J.; Formal Analysis, G.J.; Investigation, G.J.; Resources, G.J., R.S. and M.M.; Writing – Original Draft, G.J.; Writing – Review and Editing, G.J., J.I. and H.H.; Visualization, G.J.; Supervision, G.J. and J.I.; Funding Acquisition, G.J. and J.I.

